# Structural covariance networks across the lifespan, from 6-94 years of age

**DOI:** 10.1101/090233

**Authors:** Elizabeth DuPre, R. Nathan Spreng

## Abstract

Structural covariance examines covariation of grey matter morphology between brain regions and across individuals. Despite significant interest in the influence of age on structural covariance patterns, no study to date has provided a complete lifespan perspective—bridging childhood with early, middle, and late adulthood—on the development of structural covariance networks. Here, we investigate the lifespan trajectories of structural covariance in six canonical neurocognitive networks: default, dorsal attention, frontoparietal control, somatomotor, ventral attention, and visual. By combining data from five open access data sources, we examine the structural covariance trajectories of these networks from 6-94 years of age in a sample of 1580 participants. Using partial least squares, we show that structural covariance patterns across the lifespan exhibit two significant, age-dependent trends. The first trend is a stable pattern whose integrity declines over the lifespan. The second trend is an inverted-U that differentiates young adulthood from other age groups. Hub regions, including posterior cingulate cortex and anterior insula, appear particularly influential in the expression of this second age-dependent trend. Overall, our results suggest that structural covariance provides a reliable definition of neurocognitive networks across the lifespan and reveal both shared and network-specific trajectories.

## 1. Introduction

The human cerebral cortex is hierarchically organized into complex brain networks that can be considered at multiple levels of analysis (Mesulam, 1998). One such level is structural covariance, or how interindividual differences in regional brain structure covary with other brain structures across the population (Mechelli, Friston, Frackowiak, & Price, 2005; Alexander-Bloch, Giedd, & Bullmore, 2013). Structural covariance networks reflect shared variation in grey matter morphology (Mechelli et al., 2005) and are assessed using measures such as cortical thickness and regional volume. These networks exhibit reproducible organization at both a population (Alexander-Bloch et al., 2013) and individual (Tijms, Seris, Willshaw, & Lawrie, 2012) level and have been identified across species (Pagani, Bifone, & Gozzi, 2016), underscoring their role as an intrinsic feature of cortical organization. Despite this reliability, the source of grey matter shared covariance patterns is unclear and has been hypothesized to reflect both genetic and plastic influences including maturational timing (Alexander-Bloch, Raznahan, Bullmore, & Giedd, 2013).

Age is a significant moderator of both anatomical (Collin & van den Heuvel, 2013; Hagmann et al., 2010) as well as functional (Dosenbach et al., 2010; Chan, Park, Savalia, Petersen, & Wig, 2014) connectivity. Some of the most extensive age effects occur in grey matter (Giorgio et al., 2010). Grey matter organization undergoes significant structural change with age including synaptic proliferation, pruning, and eventual atrophy (Low & Cheng, 2006; Fjell et al., 2010). Normative grey matter changes do not occur simultaneously, however, and show variation across cortex (Krongold, Cooper, & Bray, 2017; Raz et al., 2005), yielding significant differences in age-related trajectories across structural covariance networks. There has therefore been substantial interest in the impacts of age on structural covariance networks, and how these age-related trajectories may differ across neurocognitive networks.

Investigations of structural covariance trajectories have largely focused on specific developmental periods including childhood and adolescence (Zielinski et al., 2010) or aging (Montembeault et al., 2012). These studies have suggested the emergence of increasing long-range structural covariance across early development (Zielinski, Gennatas, Zhou, & Seeley, 2010) and increased local covariance with advancing age (Montembeault et al., 2012). Importantly, examining structural covariance networks in isolated developmental periods may limit our understanding of the normative life cycle of each of these networks (Zuo et al., 2017). Initial work examining trajectories over multiple developmental periods has found significant inter-network variation (Hafkemeijer et al., 2014).

There has also been increasing interest in examining structural covariance networks from a lifespan perspective. However, to date existing lifespan structural covariance studies (i.e., those spanning a minimum of 35 years of development; Zuo et al., 2017) have only included subjects with a minimum age of 18 years and considered differences between young-, middle-, or older-adult groups (Li et al., 2013; Wu et al., 2012). Results from these studies have largely been in agreement with those of individual developmental periods, with distributed structural covariance shifting to more local topology in older adulthood, though the timing of this transition is unclear and has differed between middle- (Wu et al., 2012) and younger- (Li et al., 2013) adulthood. Studies have also shown differences in structural covariance trajectories by network, with primary sensory and motor networks showing few to no age-related changes across adulthood, while neurocognitive and semantic networks show a general shift from distributed to local covariance (Li et al., 2013).

Despite this significant progress in understanding structural covariance during development and aging, the authors are unaware of any studies that have examined the development of large-scale structural covariance networks across the entire lifespan, including childhood and adolescence to old age. The changes seen in the developmental trajectory of large-scale functional networks (Zuo et al., 2017) suggest that a lifespan study of structural covariance networks may provide an important complement, yielding insights into cortical organization at the level of grey matter morphology. Indeed, previous work by Zielinski and colleagues (2010) supports the reflection of a network’s functional specialization in its age-specific structural covariance pattern. Based on previous findings, we hypothesized that the distributed neurocognitive default, dorsal attention, frontoparietal control and ventral attention networks would show an inverted U-shaped trajectory of increasingly distributed structural covariance in early development, before shifting to more local covariance in advanced aging. Following results reported by Li and colleagues (2013) of age-independent patterns of structural covariance in somatomotor and visual networks across the adult lifespan, we predicted no age-dependent patterns of structural covariance in these networks. To examine these hypothesized lifespan trajectories, whole-brain structural covariance was assessed in a seed-based multivariate analysis (Spreng & Turner, 2013; Persson et al., 2014). This seed-based multivariate investigation allowed for the data-driven identification of significant age-related trajectories, based on the structural covariance of cortical grey matter with the chosen seed regions. We examine trajectories of structural covariance networks across the lifespan to consider what these changes might reveal about their developmental organization.

## 2. Materials and Methods

In this study, our primary aim was to provide comprehensive mapping of the neurocognitive large-scale structural covariance networks across the entire lifespan. We collapsed cross-sectional data across five publicly available datasets to provide a normative sample ranging from six to ninety-four years of age. This also afforded us sufficient power for reliable estimates of structural covariance networks at six developmental epochs: Age Group 1 (6-15y), Age Group 2 (16-25y), Age Group 3 (26-35y), Age Group 4 (36-59y), Age Group 5 (60-75y), and Age Group 6 (76-94y). We assessed the structural covariance of six large-scale neurocognitive networks well represented in the literature: The default network (DN), dorsal attention network (DAN), frontoparietal control network (FPCN), somatomotor network (SM), ventral attention network (VAN), and visual systems.

### 2.1. Image Acquisition

Data were collated from five open access data sources: National Institutes of Health Pediatric MRI Data Repository (NIH-Peds; Brain Development Cooperative Group & Evans, 2007): Release 5, Human Connectome Project (HCP): 500 subjects release, Nathan Kline Institute-Rockland Sample (NKI-RS; Nooner et al., 2012): Release 5, Open Access Series of Imaging Studies (OASIS), and Alzheimer’s Disease Neuroimaging Initiative (ADNI). A complete listing of T1-weighted anatomical image acquisition procedures for each data source is provided in Table 1.

**Table 1.** Image Acquisition Parameters by Data Source.

|  | Scanner<br>Strength<br>(T) | TR<br>(ms) | TE<br>(ms) | TI<br>(ms) | TD<br>(ms) | Flip<br>Angle<br>(deg.) | FOV<br>(mm) | Voxel Size<br>(mm) |
| --- | --- | --- | --- | --- | --- | --- | --- | --- |
| NIH-Peds | 1.5 | 22-25 | 10-11 | | | | 256×160-180 | $1.0 \times 1.0 \times 1.0-1.5$ |
| HCP | 3 | 2400 | 2.14 | 1000 | 0 | 8 | 224×224 | $0.7 \times 0.7 \times 0.7$ |
| NKI-RS | 3 | 1900 | 2.52 | 900 | 0 | 9 | 250×250 | $1.0 \times 1.0 \times 1.0$ |
| OASIS | 1.5 | 9.7 | 4.0 | 20 | 200 | 10 | 256×256 | $1.0 \times 1.0 \times 1.25$ |
| ADNI | 1.5 | 2400 | min.<br>full | 1000 | 0 | 8 | 240×240 | $0.94 \times 0.94 \times 1.2$ |

### 2.2. Participant Characteristics

From each sample, only healthy control participants older than six years of age with no diagnosed history of DSM Axis I or II disorder were considered. Six years was chosen as the lowest estimate for lifespan characterization, since previous work has indicated that normalization for children less than six years is likely to introduce significant artifacts (Muzik, Chugani, Juhász, Shen, & Chugani, 2000) as grey matter volume in younger children is less than 95% of that observed in adults (Caviness, Kennedy, Richelme, Rademacher, & Filipek, 1996). For individuals meeting these criteria, the T1-weighted anatomical image was selected. In the case of longitudinal data, only the first time point was selected for each participant.

All T1-weighted anatomical images (n = 1667) were visually inspected for quality assurance: images that showed evidence of artifacts were excluded (n = 87), yielding a final sample size of n = 1580 (age *M* = 35y, *SD* = 23y, Range = 6 - 94y; 659 males; 859 scanned at 1.5T and 721 at 3T). Participants were then sorted into the following age groups: Age Group 1 (6-15y), Age Group 2 (16-25y), Age Group 3 (26-35y), Age Group 4 (36-59y), Age Group 5 (60-75y), and Age Group 6 (76-94y). See Table 2 for sample sizes and participant characteristics by age group.

**Table 2.** Participant Characteristics by Age Group.

| Age Group | Sample Size<br>(Males) | Age in Years<br>$M (SD)$ | Scanned at<br>1.5T/3T |
| --- | --- | --- | --- |
| 6-15y | 330 (159) | 10 (2.66) | 306/24 |
| 16-25y | 302 (139) | 21 (2.8) | 176/126 |
| 26-35y | 472 (192) | 30 (2.74) | 31/441 |
| 36-59y | 139 (38) | 49 (6.29) | 68/71 |
| 60-75y | 203 (82) | 70 (4.16) | 157/46 |
| 76-94y | 134 (49) | 81 (4.28) | 121/13 |

### 2.3. Segmentation and Preprocessing

Each age group was separately submitted to voxel-based morphometry (Ashburner & Friston, 2000) using the VBM8 toolbox (www.neuro.uni-jena.de/vbm/) implemented in Matlab (MATLAB 8.0, MathWorks, Natick, MA, 2012). Images were first segmented into grey matter, white matter, and cerebrospinal fluid using an extension of the New Segmentation algorithm. Grey matter images for this age group were then affine registered to the MNI template and carried to the Diffeomorphic Anatomical Registration through Exponentiated Lie Algebra toolbox (DARTEL; Ashburner, 2007) where they were iteratively aligned to create an age-group-specific template in MNI space. The six resulting age-group-specific templates were themselves then iteratively aligned again using DARTEL to create a study-specific template in MNI space. Importantly, this study-specific template equally weighted each of the age ranges represented by the six age groups.

Finally, previously segmented images were aligned to the study-specific template of interest using DARTEL high-dimensional normalization within VBM8. Non-linear only modulation was applied to grey matter images to derive regional differences in grey matter volume, correcting for total intracranial volume. Modulated grey matter images were iteratively smoothed to 8mm FWHM using *3dBlurToFWHM* in AFNI (Cox, 1996) and carried forward for further analysis.

### 2.4. Network Identification

In this study, we sought to examine the structural covariance of the large-scale neurocognitive networks, including the DN, DAN, FPCN, SM, VAN, and visual networks. To examine each of these six networks, grey matter volumes for selected high-confidence seeds reported in Yeo et al. (2011) were extracted. Although Yeo and colleagues (2011) report high-confidence seeds for seven networks, we chose to exclude the reported “limbic network” as recent work has raised concerns regarding its test-retest reliability (Holmes et al., 2015).

For each of the six remaining networks we selected the top two high-confidence seeds reported by Yeo and colleagues (2011) as well as the contralateral seed regions, where contralateral seeds were chosen by changing the sign of the x-coordinate on each of the original high-confidence seeds. An exception to this procedure was made for the DN, which is known to separate into anterior and posterior components (Uddin, Kelly, Biswal, Castellanos, & Milham, 2009). Therefore, for the DN we selected the highest confidence seed and its contralateral seed. We then selected the second highest confidence seed in posterior cingulate cortex as well as the fourth highest confidence seed in medial prefrontal cortex, in order to ensure that both the anterior and posterior DN components were represented in structural covariance estimates. A listing of all 24 seeds for the six networks examined is presented in Table 3 along with their respective anatomical label, and a visual representation of their location on cortex is presented in Figure 1.

**Table 3.** Selected Seeds for Each Network.

| Network Affiliation | $x$ | $y$ | $z$ | Laterality | Anatomical Label |
| --- | --- | --- | --- | --- | --- |
| Default | -7 | 49 | 18 | L | Medial Prefrontal Cortex |
|  | -7 | -52 | 26 | L | Posterior Cingulate Cortex |
|  | -41 | -60 | 29 | L | Inferior Parietal Lobule |
|  | 41 | -60 | 29 | R | Inferior Parietal Lobule |
| Dorsal Attention | -22 | -8 | 54 | L | Frontal Eye Fields |
|  | 22 | -8 | 54 | R | Frontal Eye Fields |
|  | -51 | -64 | -2 | L | Middle Temporal Motion Complex |
|  | 51 | -64 | -2 | R | Middle Temporal Motion Complex |
| Frontoparietal Control | -40 | 50 | 7 | L | Frontal Pole |
|  | 40 | 50 | 7 | R | Frontal Pole |
|  | -43 | -50 | 46 | L | Anterior Inferior Parietal Lobule |
|  | 43 | -50 | 46 | R | Anterior Inferior Parietal Lobule |
| Ventral Attention | -5 | 15 | 32 | L | Anterior Cingulate Cortex |
|  | 5 | 15 | 32 | R | Anterior Cingulate Cortex |
|  | -31 | 11 | 8 | L | Anterior Insula |
|  | 31 | 11 | 8 | R | Anterior Insula |
| Somatomotor | -41 | -20 | 62 | L | Precentral Gyrus (Hand) |
|  | 41 | -20 | 62 | R | Precentral Gyrus (Hand) |
|  | -55 | -4 | 26 | L | Precentral Gyrus (Tongue) |
|  | 55 | -4 | 26 | R | Precentral Gyrus (Tongue) |
| Visual | -3 | -74 | 23 | L | Extrastriate Visual Cortex |
|  | 3 | -74 | 23 | R | Extrastriate Visual Cortex |
|  | -16 | -74 | 7 | L | Visual Area 1 |
|  | 16 | -74 | 7 | R | Visual Area 1 |

Coordinates ( $x$ , $y$ , $z$ ) are in MNI stereotaxic space.

**Figure 1.**
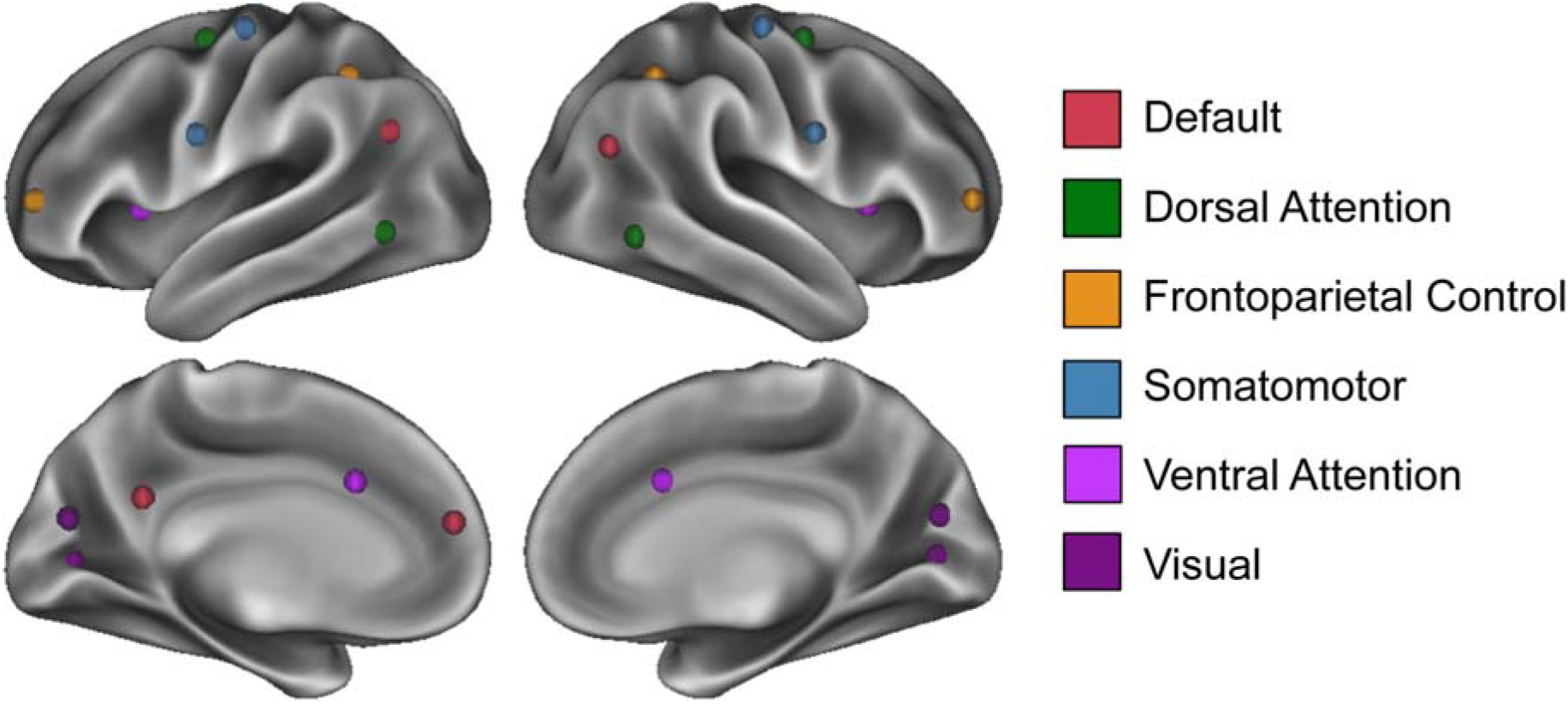
Selected seeded regions. The four selected seeded regions for each of the six neurocognitive networks are depicted in colors corresponding to their Yeo and colleagues (2011) labeling.

For each seed, grey matter volumes were extracted from a 10.5mm edge cubical ROI. Extracted grey matter volumes were then averaged across the four seeds for each participant. We chose to average grey matter volumes from multiple seeds to provide reliable, long-range estimates of network-specific structural covariance. This is in contrast to the more local estimates of structural covariance provided by grey matter volume from a singe seed region. All six neurocognitive networks were examined by averaging the extracted grey matter volumes for each of the network-specific seeds, resulting in a 1580 × 1 vector for each network. For each of the analyses, this vector **Y** represented the average grey matter volume for each participant of key nodes within the network. The resulting **Y** vectors were submitted to Partial Least Squares (PLS; McIntosh, Bookstein, Haxby, & Grady, 1996). Additionally submitted to PLS were matrices of participant structural images, **X**, where **X** is an *N* subjects x *N* voxels matrix representing voxel-wise estimates of grey matter volume for each participant.

### 2.5. Partial Least Squares (PLS) Analyses

PLS is a data-driven multivariate statistical technique capable of identifying patterns of structural covariance (Spreng & Turner, 2013; Persson et al., 2014). We utilized seed PLS to identify patterns of covariance between grey matter integrity in seed regions and whole brain structural MRI images (for a review, see Krishnan, Williams, McIntosh, & Abdi, 2011). Here, we adopt the nomenclature used in Mišić and colleagues (2016).

#### 2.5.1. Derivation of Covariance Matrix

For experimental analyses, our seed value was the average grey matter volume of four selected high-confidence seeds reported in Yeo et al. (2011). The vector **Y** representing this average grey matter volume was cross-correlated with a matrix **X** of participant’s structural images. Importantly, this participant image matrix contained six sub-matrices **X**_1..6_ corresponding to each age group. We retained this age group organization in our PLS analyses in order to directly compare age groups in their structural covariance between average network and whole brain grey matter volume. The vector **Y** can therefore be considered as containing six sub-vectors, corresponding to the participant age groups. Both the grey matter volume vector and image matrix were centered and normalized within age groups such that their cross-correlation resulted in a covariance vector **Z** according to:

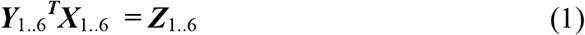

Note that this covariance vector is equivalent to a correlation vector due to the described within-group normalization. The resulting covariance vector **Z** measures the degree to which the network average and whole brain grey matter volumes covary at a voxel-wise level across participants.

#### 2.5.2. Singular Value Decomposition

Using singular value decomposition (SVD; Eckart & Young, 1936), the covariance vector **Z** from Eq. (1) was then decomposed into:

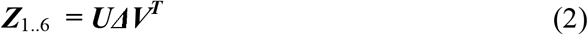

where **V** is the orthonormal matrix of right singular vectors, **U** is the orthonormal matrix of left singular vectors, and Δ is the diagonal matrix of singular values. The right and left singular vectors represent the grey matter seed integrity profiles and spatial patterns that best characterize the covariance vector **Z**. The triplet of the right and left singular vectors and the singular values forms a set of mutually orthogonal Latent Variables (LVs) where the number of LVs derived is equal to the rank of the covariance vector **Z**. In our analyses, this identified six LVs for each network corresponding to the six sub-matrices of **Z**. Each LV was tested for statistical significance with 5000 permutations and cross-validated for reliability with 1000 bootstraps. Bootstrap ratios, derived from dividing the weight of the singular-vector by the bootstrapped standard error, are equivalent to z-scores and were used to threshold significant LV spatial patterns at a 95% confidence interval for projection and interpretation.

Patterns were considered for further analysis based on two criteria. First, LVs must be statistically significant by permutation testing at the level of *p* < 0.001. Second, LVs must account for a minimum of 5% of the covariance in the data.

#### 2.5.3. Derivation of Subject Scores

We also quantified individual contributions to each LV by deriving subject scores. Of particular interest in this work are the subject scores known in PLS nomenclature as “brain scores,” which assess the contribution of each individual to the group structural covariance pattern. Multiplying the original matrix **X**_1..6_ of participant structural images by the matrix of right singular vectors **V** derive these brain scores as follows:

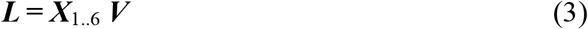

where **L** is a matrix of brain scores. Recall from Eq. (2) that the right singular vector **V** represents the seed-integrity profiles that best characterizes the covariance matrix **Z**, such that multiplying this singular vector by the participant structural images derives the seed integrity profiles for each participant that reflect their contribution to the group structural covariance pattern. The matrix of brain scores **L** was extracted for each LV where, for each participant, this brain score value represents a weighted value of grey matter integrity within the regions identified in the group image.

By correlating brain score values for all subjects within each of the six age groups with their input grey matter integrity values, we were able to assess grey matter integrity in these regions for each age group separately. Computed confidence intervals on these correlations provide a means to assess the reliability of the structural covariance patterns in each age group; confidence intervals which cross zero are considered unreliable and are not interpreted in the results. To account for potential confounds, we ran a multiple regression of these brain scores controlling for scanner strength and gender. Although we present results corrected for age and gender, controlling for these variables did not qualitatively affect the results (see Supplementary Figure 1 for an exemplar network). Corrected brain scores were plotted against age to visualize the covariance of the associated spatial pattern across the population. Due to the heterogeneity of resulting age-dependent trajectories, summary statistics for models fit to these corrected brain scores are available in Supplementary Tables 1 and 2. For those models who show a “peak” in their age-dependent trajectory, the age at which this functional maximum occurs is noted in Supplementary Table 2.

## 3. Results

We investigated the structural covariance of previously identified large-scale neurocognitive networks including the DN, DAN, FPCN, SM, VAN, and visual networks. Using PLS, we identified patterns of structural covariance for each of the six networks examined.

### 3.1. Neurocognitive Network Structural Covariance Patterns

PLS analyses of each of the large-scale networks examined yielded multiple significant latent variables (LVs), corresponding to reliable patterns of structural covariance within each network. We review significant results for each of the networks in turn.

#### 3.1.1. Default Network

Two significant LVs were identified for the DN and are presented in Figure 2. In the first LV (*p* < 0.0002; 61.57% covariance explained), seeded regions, along with homologous contralateral regions, covary together as well as with parahippocampal cortex and lateral temporal cortex (Fig 2A). Covariance extended to non-canonical DN regions including posterior insula. All age groups showed a robust positive association with this pattern (Figure 2B); this suggests that this latent variable corresponds to the structural covariance of the DN as it is preserved across the lifespan. Extracted brain scores (Fig 2C) revealed that the integrity of this structural covariance pattern declines with advancing age rapidly before reaching a plateau at approximately 70 years of age.

**Figure 2.**
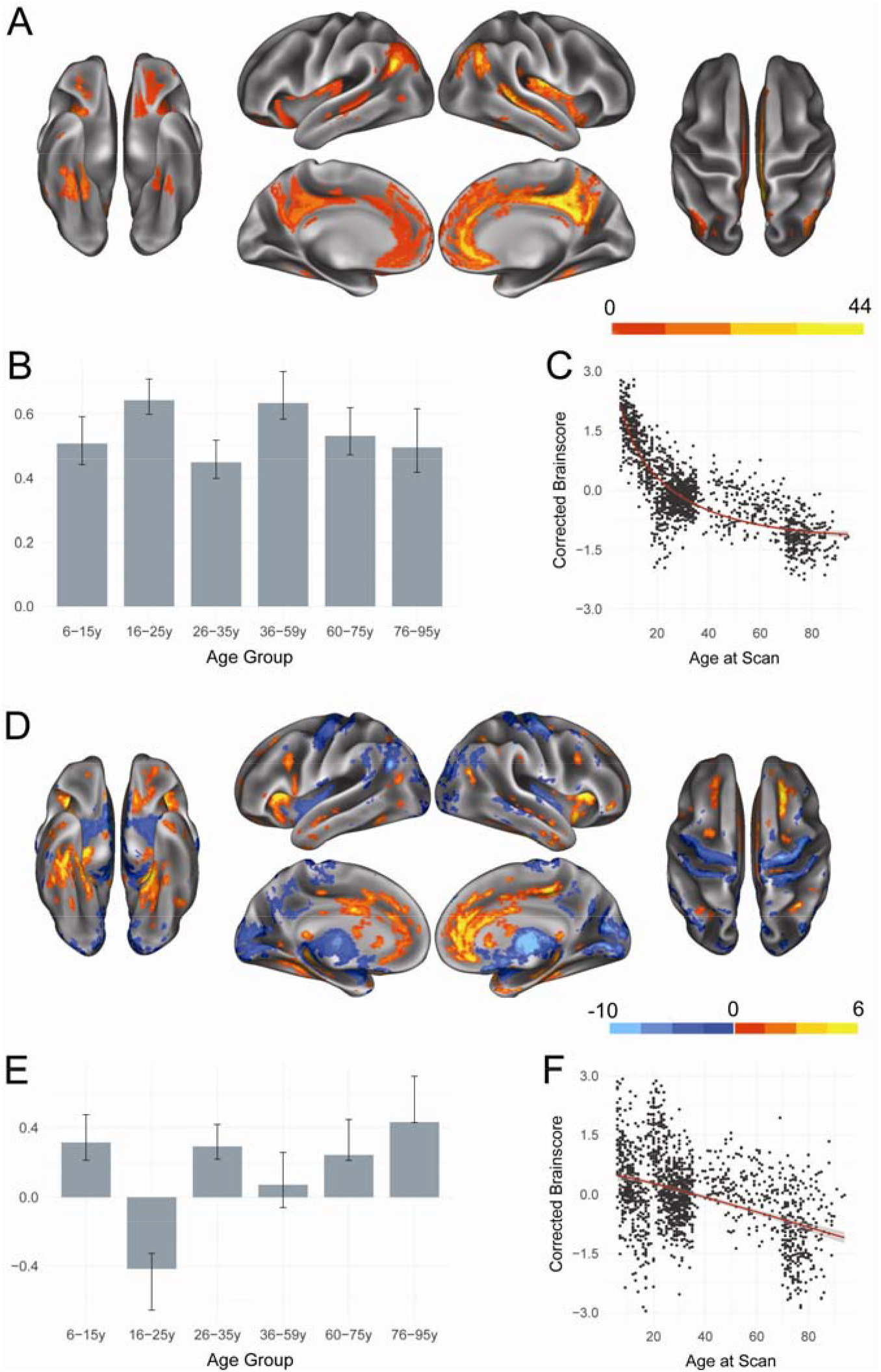
Structural covariance of the default network. (A) The spatial pattern for the first latent variable, thresholded at 95% of the bootstrap ratio. (B) The bootstrapped correlation of brain scores with the averaged grey matter volume estimates of default network seeds by age group for the first latent variable. (C) The individual brain scores from the first latent variable corrected for scanner strength and gender are plotted as a function of age. (D) Spatial pattern for the second latent variable, thresholded at 95% of the bootstrap ratio. (E) The bootstrapped correlation of brain scores with averaged grey matter volume estimates of default network seeds by age group for the second latent variable. (F) The individual brain scores from the second latent variable corrected for scanner strength and gender are plotted as a function of age.

The second significant LV (*p* < 0.0002; 13.71% covariance explained) showed structural covariance patterns of developmental change in the DN. Age Group 2 (16-25y) showed a unique pattern of increased structural covariance with medial prefrontal cortex and anterior insula compared to all other age groups examined (Figure 2D, E). Age groups with reliable correlations of brain scores and behavior—those for which the confidence interval did not cross zero and were therefore considered interpretable—included the Age Group 1 (6-15y), Age Group 3 (26-35y), Age Group 5 (60-75y), and Age Group 6 (76-94y) cohorts. Compared to Age Group 2 (16-25y), each of these cohorts showed relatively increased structural covariance between seeded DN regions and sensorimotor structures including motor and visual cortices as well as thalamus. Across the lifespan, this pattern shows a nearly linear decrease with advancing age (Figure 2F), suggesting that older adults are less strongly aligning to the structural covariance pattern depicted in Figure 2D.

#### 3.1.2. Dorsal Attention Network

Two significant LVs were identified for the DAN and are presented in Figure 3. In line with results presented for the DN, the first significant DAN LV (*p* < 0.0002; 70.93% covariance explained) showed seeded regions positively covarying together as well as with canonical DAN regions such as intraparietal sulcus (Figure 3A). Covariance also extended to other, non-canonical DAN regions, including posterior insula and subgenual cingulate. All age groups showed a robust association with this pattern (Figure 6B). Brain scores reveal that the integrity of this pattern shows rapid decline with advancing age before plateauing at approximately 70 years (Figure 3C).

**Figure 3.**
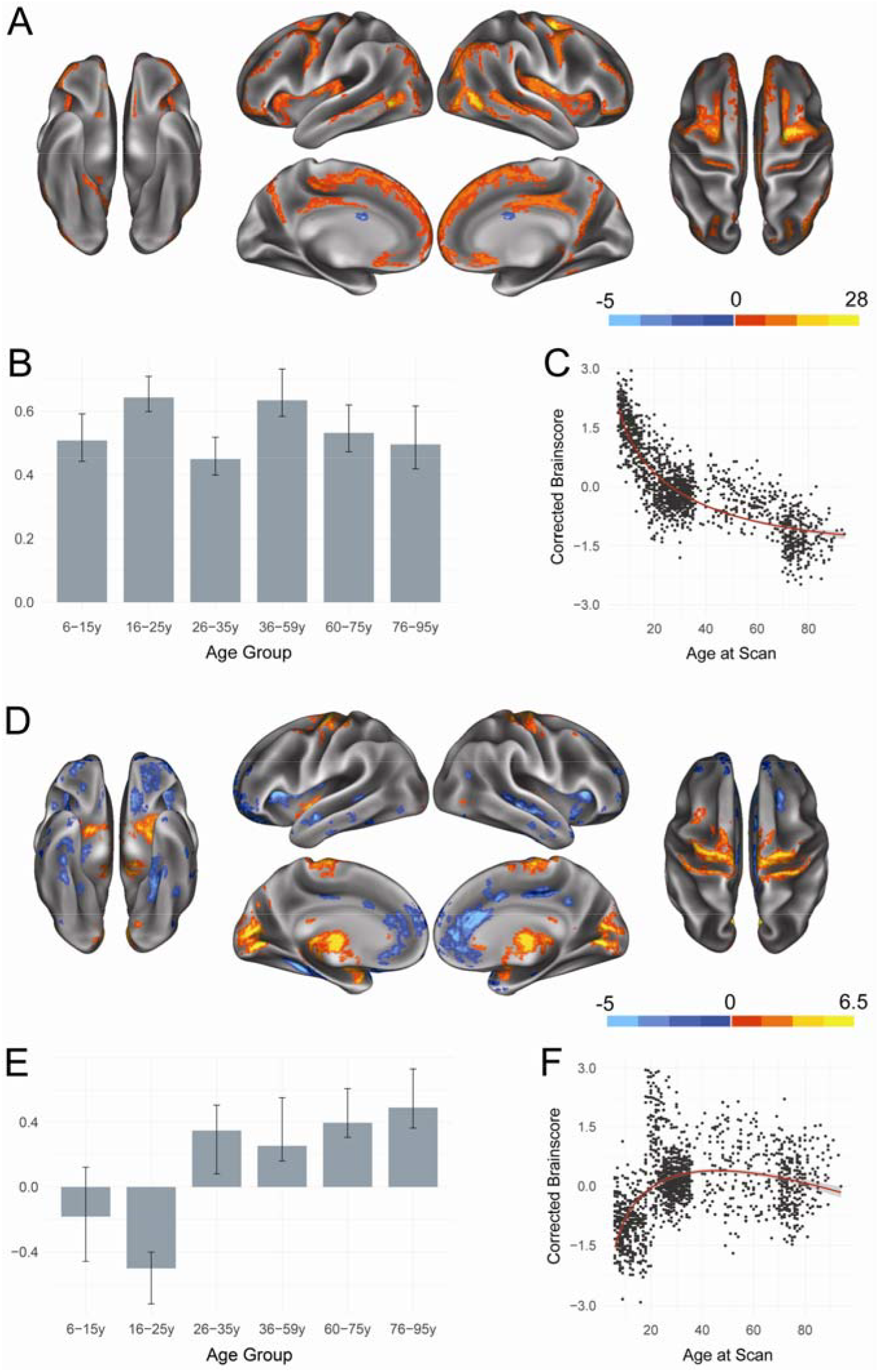
Structural covariance of the dorsal attention network. (A) The spatial pattern for the first latent variable, thresholded at 95% of the bootstrap ratio. (B) The bootstrapped correlation of brain scores with averaged grey matter volume estimates of dorsal attention network seeds by age group for the first latent variable. (C) The individual brain scores from the first latent variable corrected for scanner strength and gender are plotted as a function of age. (D) Spatial pattern for the second latent variable, thresholded at 95% of the bootstrap ratio. (E) The bootstrapped correlation of brain scores with averaged grey matter volume estimates of dorsal attention network seeds for the second latent variable. (F) The individual brain scores from the second latent variable corrected for scanner strength and gender are plotted as a function of age.

The second significant LV (*p* < 0.0002; 9.52% covariance explained) revealed developmental changes in the structural covariance pattern of the DAN. Age Group 2 (16-25y) showed uniquely increased structural covariance with medial prefrontal cortex and anterior insula. Older age groups show relatively increased structural covariance between the seeded DAN regions and areas including motor and visual cortices as well as subcortical structures. Inspection of brain scores (Figure 3F) reveals an inverted U-shaped trajectory, with integrity of the structural covariance pattern reaching its peak in middle adulthood, while very young and very old individuals show significantly less integrity for the derived group structural covariance patterns.

#### 3.1.3. Frontoparietal Control Network

Two significant LVs were identified for the FPCN and are depicted in Figure 4. Similar to results seen for the DN and DAN, the first significant LV (*p* < 0.0002; 78.05% covariance explained) showed a structural covariance pattern that was positively associated with all examined age groups, but showed a non-linear decline in integrity across the lifespan. Seeded FPCN regions positively covary together, as well as with structures consistently associated with cognitive control, such as lateral prefrontal cortex, and non-canonical FPCN regions, such as posterior insula.

**Figure 4.**
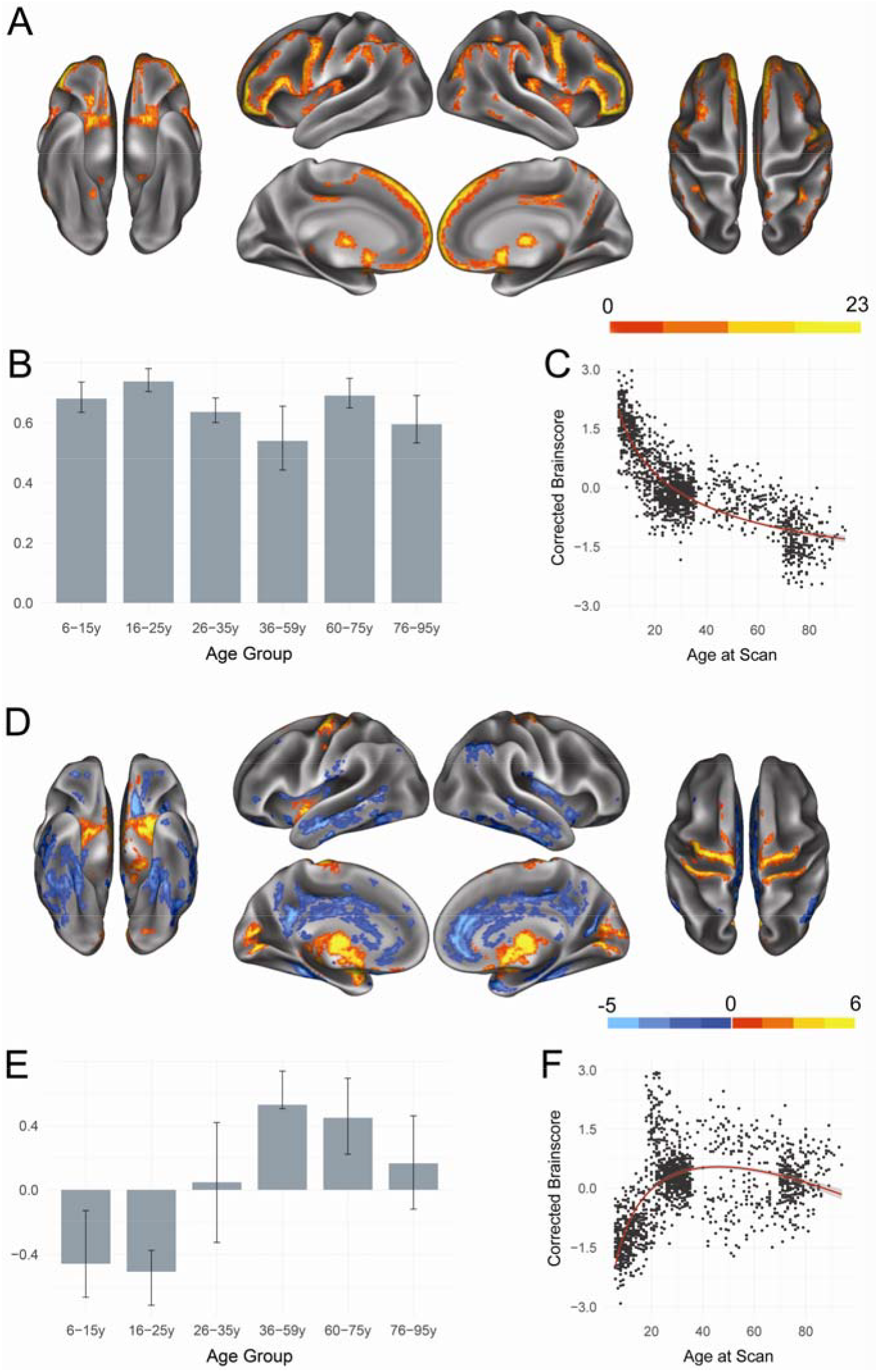
Structural covariance of the frontoparietal control network. (A) The spatial pattern for the first latent variable, thresholded at 95% of the bootstrap ratio. (B) The bootstrapped correlation of brain scores with averaged grey matter volume estimates of frontoparietal control network seeds. (C) The individual brain scores from the first latent variable corrected for scanner strength and gender are plotted as a function of age. (D) Spatial pattern for the second latent variable, thresholded at 95% of the bootstrap ratio. (E) The bootstrapped correlation of brain scores with averaged grey matter volume estimates of default network seeds by age group for the second latent variable. (F) The individual brain scores from the second latent variable corrected for scanner strength and gender are plotted as a function of age.

The second LV (*p* < 0.0002; 7.96% covariance explained) revealed developmental trajectories of structural covariance patterns in the FPCN. There was a significant dissociation between Age Groups 1 and 2 (6-25y) as compared to middle and late Age Groups 4 and 5 (36-75y). Younger age groups show increased structural covariance with structures both within the canonical FPCN such as precuneus as well as with non-canonical regions such as lateral temporal cortex. Older age groups, however, show relatively increased structural covariance for sensorimotor structures such as motor cortex and thalamus. Brain scores suggest an inverted U-shaped trajectory similar to that seen for the DAN, with the integrity of the structural covariance pattern at its highest levels in middle adulthood.

#### 3.1.4. Somatomotor Network

Two significant LVs were identified for the SM and are depicted in Figure 5. In agreement with the previously reported networks, the first significant LV (*p* < 0.0002; 72.91% covariance explained) showed a structural covariance pattern that is positively associated with all examined age groups and shows a non-linear decline with advancing age. Seeded regions covaried together as well as with the motor strip. Covariance extended to areas outside the canonical motor network such as lateral prefrontal cortex and subcortical regions.

**Figure 5.**
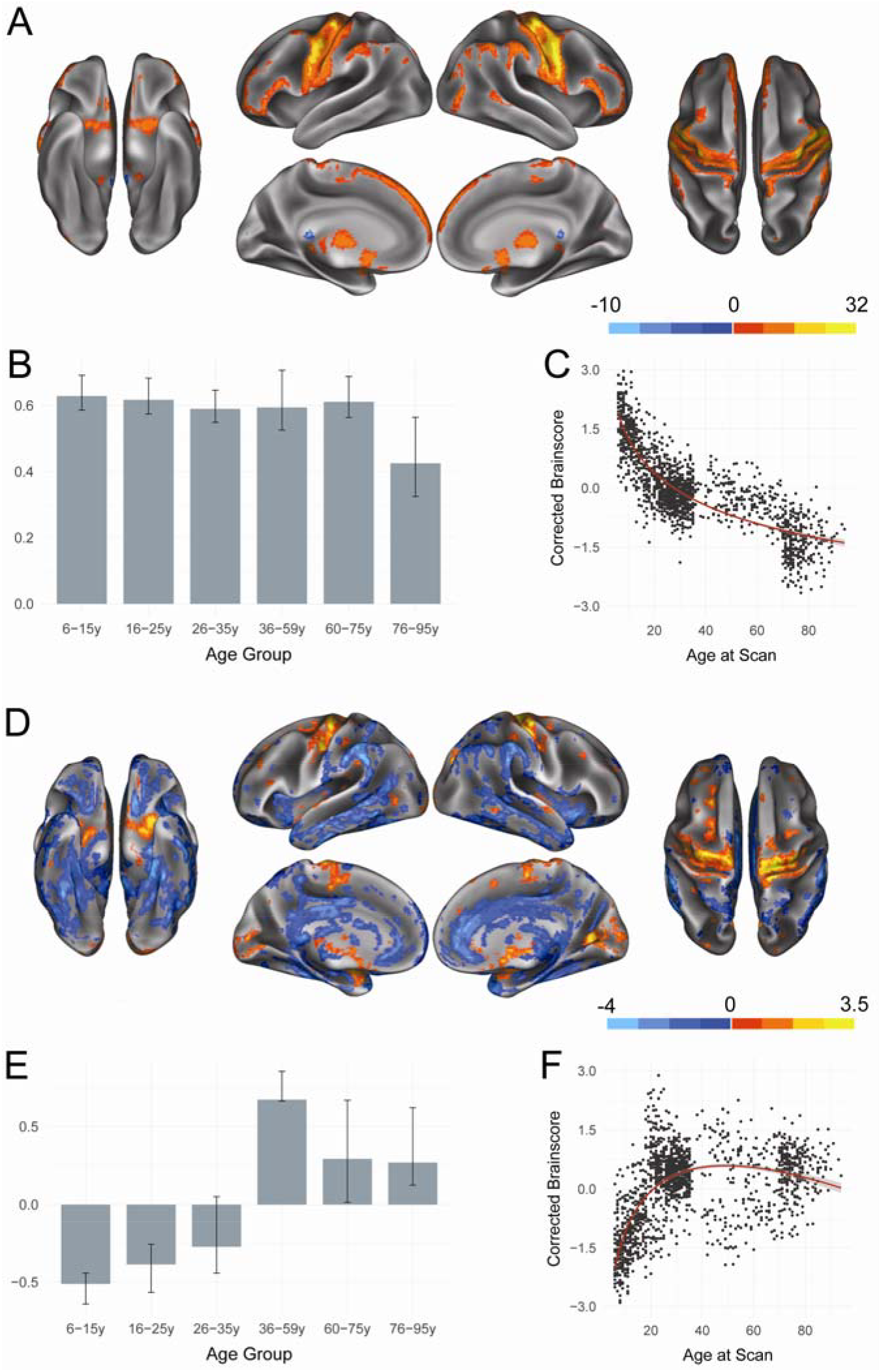
Structural covariance of the somatomotor network. (A) The spatial pattern for the first latent variable, thresholded at 95% of the bootstrap ratio. (B) The bootstrapped correlation of brain scores with bootstrapped averaged grey matter volume estimates of somatomotor network seeds by age group for the first latent variable. (C) The individual brain scores from the first latent variable corrected for scanner strength and gender are plotted as a function of age. (D) Spatial pattern for the second latent variable, thresholded at 95% of the bootstrap ratio. (E) The bootstrapped correlation of brain scores with averaged grey matter volume estimates of somatomotor network seeds by age group for the second latent variable. (F) The individual brain scores from the second latent variable corrected for scanner strength and gender are plotted as a function of age.

The second LV (*p* < 0.0002; 9.17% covariance explained) showed a significant dissociation between Age Groups 1 and 2 (6-25y) as compared to Age Groups 4, 5, and 6 (36-94y). Younger age groups show increased structural covariance with structures outside of the canonical motor network such as lateral temporal cortex and mid-insula, while older age groups show relatively increased structural covariance local to the seed regions and to thalamus. Similar to the individual subject score trajectories seen for the DAN and FPCN, there is an inverted U-shaped trajectory in the integrity of this structural covariance pattern, with integrity reaching a peak in middle adulthood before beginning to decline.

#### 3.1.5. Ventral Attention Network

Two significant LVs were identified for the VAN and are presented in Figure 6. The first LV (*p* < 0.0002; 70.93% covariance explained) again shows a structural covariance pattern positively associated with all examined age groups. Seeded VAN regions positively covary together and with the mid-and posterior insula as well as with the medial prefrontal cortex. Extracted brain scores revealed a non-linear decline in the integrity of this structural covariance pattern across the lifespan.

**Figure 6.**
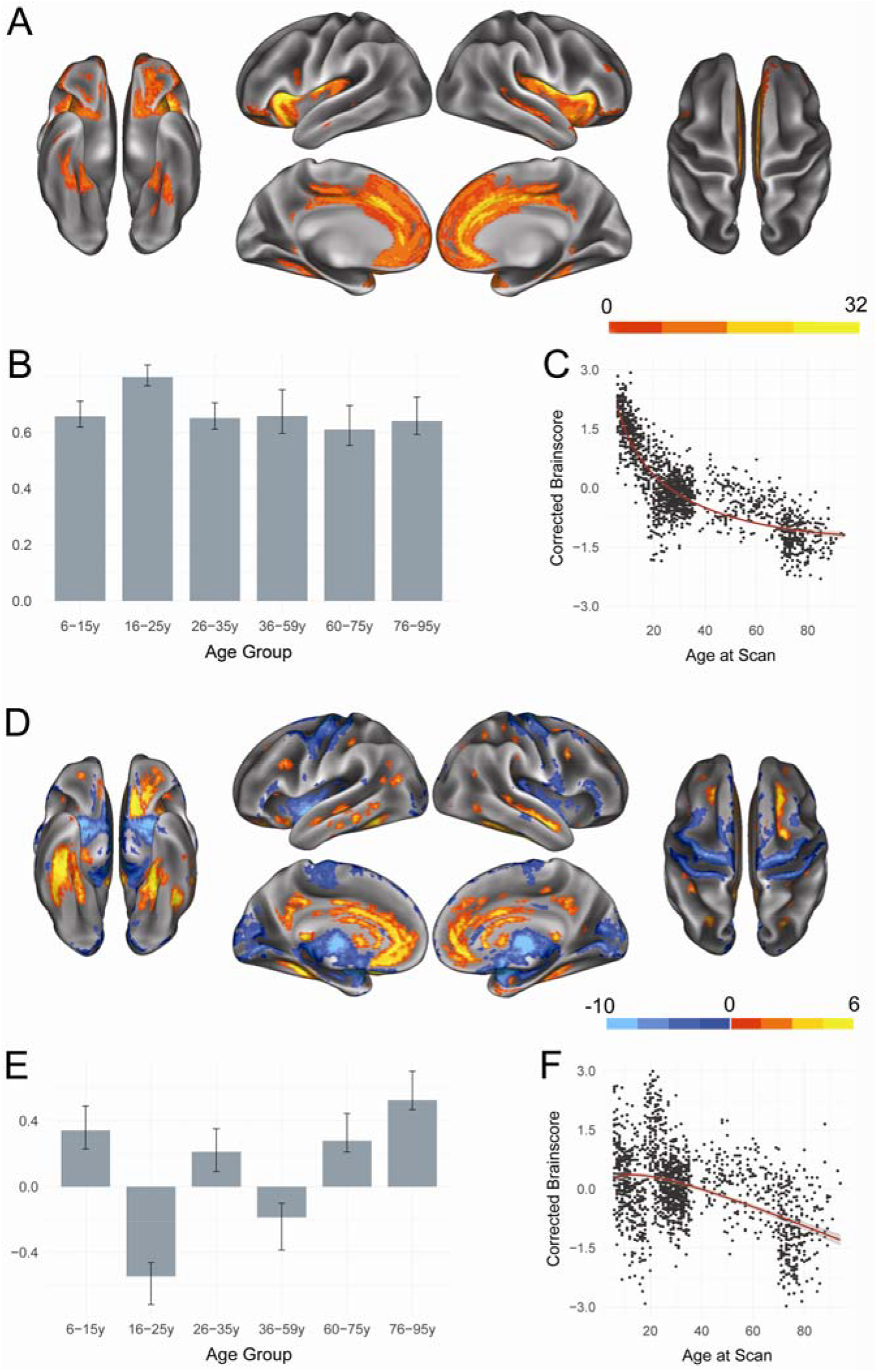
Structural covariance of the ventral attention network. (A) The spatial pattern for the first latent variable, thresholded at 95% of the bootstrap ratio. (B) The bootstrapped correlation of brain scores with averaged grey matter volume estimates of ventral attention network seeds by age group for the first latent variable. (C) The individual brain scores from the first latent variable corrected for scanner strength and gender are plotted as a function of age. (D) Spatial pattern for the second latent variable, thresholded at 95% of the bootstrap ratio. (E) The bootstrapped correlation of brain scores with averaged grey matter volume estimates of ventral attention network seeds by age group for the second latent variable. (F) The individual brain scores from the second latent variable corrected for scanner strength and gender are plotted as a function of age.

The second significant LV (*p* < 0.0002; 13.83% covariance explained) revealed a pattern of developmental change similar to that seen in DN, with individuals in Age Group 2 (16-25y) and Age Group 4 (36-59y) showing a unique structural covariance pattern compared to all other age groups. Specifically, these two groups showed increased structural covariance with medial prefrontal as well as parahippocampal cortex. Other age groups showed increased structural covariance with sensorimotor structures such as motor and visual cortices. Similarly to the DN, there is a near linear decrease in structural integrity across the lifespan, with older adults showing decreased structural covariance between seeded VAN regions and sensorimotor structures.

#### 3.1.6. Visual Network

One significant LV (*p* < 0.0002; 58.28% covariance explained) was identified for the visual network and is presented in Figure 7. As in previous networks, the significant LV revealed a structural covariance pattern that was positively associated with all examined age groups and non-linearly declined with age. Seeded visual regions showed positive structural covariance with visual cortex as well as with non-canonical visual regions such as the posterior insula and mid-cingulate.

**Figure 7.**
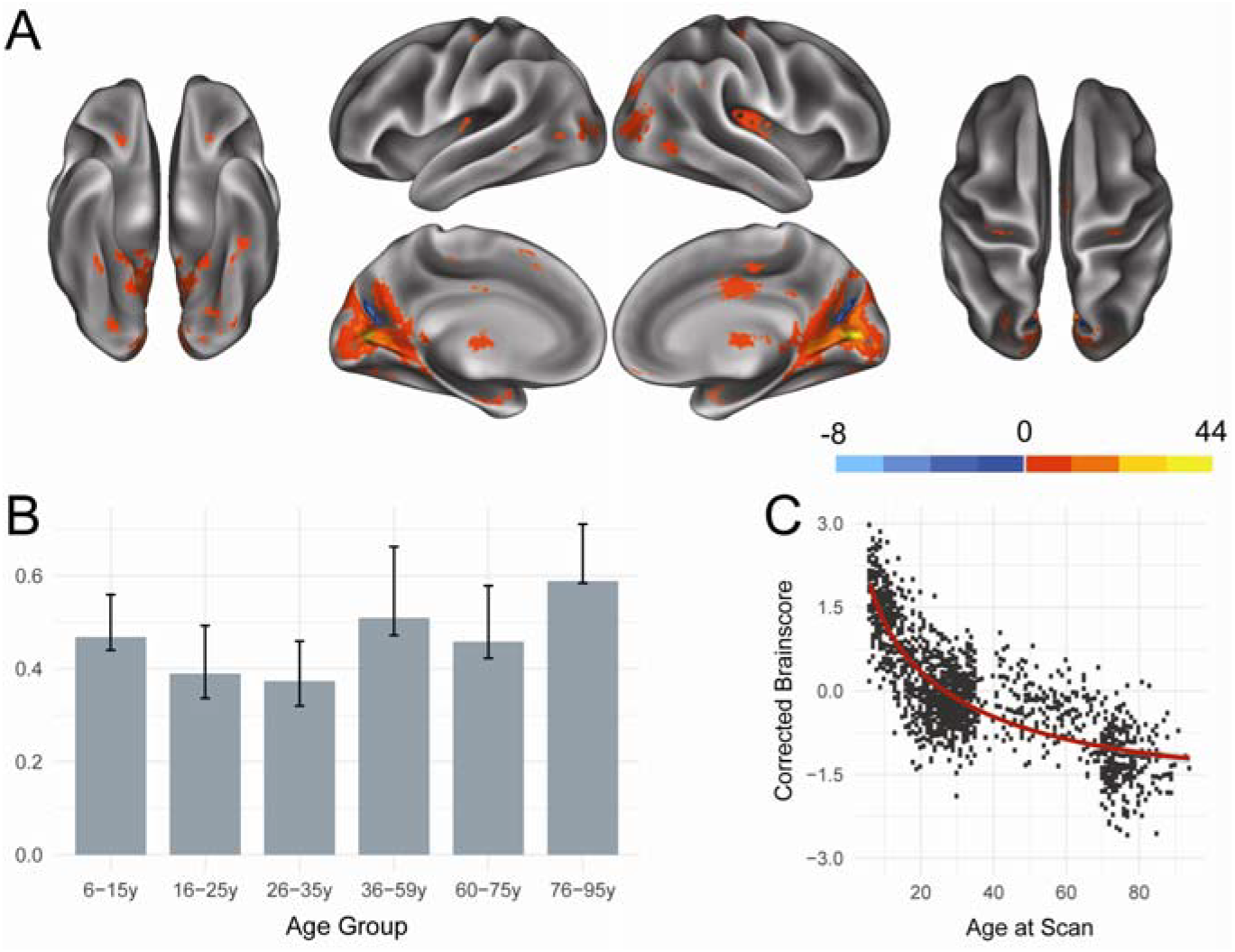
Structural covariance of the visual network. (A) The spatial pattern for the first latent variable, thresholded at 95% of the bootstrap ratio. (B) The bootstrapped correlation of brain scores with averaged grey matter volume estimates of visual network seeds by age group for the first latent variable. (C) The individual brain scores from the first latent variable corrected for scanner strength and gender are plotted as a function of age.

## 4. Discussion

In this study, we examined the lifespan trajectories of structural covariance with publicly available cross-sectional data. For the six neurocognitive networks examined, our results revealed two broad developmental patterns: a stable pattern of structural covariance that reflects network-specific features and persists across the lifespan, and an age-dependent pattern of structural covariance that reveals shared age-related trajectories of structural covariance across networks.

### 4.1 Persistent Patterns of Structural Covariance

Across all networks, the first significant latent variable identified a structural covariance pattern whose spatial extent was unique to the network of interest and persisted across age groups. Despite the stability of these structural covariance patterns over the lifespan, inspection of individual subject or “brain” scores (panel C, Figures 2-7) revealed that integrity of these patterns declines rapidly with advancing age before plateauing at approximately 70 years.

These findings extend on previous work showing a sharp decline in within-network structural covariance from young adulthood to middle age that persists into older adulthood (Li et al., 2013). Although our results show that network-specific structural covariance patterns were stable across the lifespan, we find that children and adolescents show even higher levels of integrity to these structural covariance patterns compared to young adults. This decline in integrity to structural covariance patterns over the lifespan may be related both to the increase in myelination across early development and its effects on grey-white matter tissue contrast (Lenroot & Giedd, 2006), as well as to the decline of cortical grey matter volume with age (Allen, Bruss, & Damasio, 2005).

### 4.2 Age-Dependent Patterns of Structural Covariance

In addition to stable patterns of structural covariance, DN, DAN, FPCN, SM, and VAN networks showed an additional, age-dependent pattern that differentiated young adulthood from either end of the lifespan.

Examination of brain scores in the DAN, FPCN, and SM (panel F, Figures 3-5) reveals that these align with an inverted U-shaped trajectory. These latent variables also showed overlapping features of structural covariance at a group level (panel D of Figures 3-5). In young adulthood, seeded regions showed structural covariance with areas including medial prefrontal cortex, posterior cingulate, insular cortex, and temporal cortex—association cortices corresponding to functional hubs (van den Heuvel & Sporns, 2013). In both childhood and older adulthood, however, seeded regions showed structural covariance with sensorimotor structures including motor and visual cortices and thalamus. These findings support previous work showing that structural covariance networks grow increasingly distributed over early development before shifting to a more localized topology in advanced aging (Wu et al., 2012). Our results in the DAN, FPCN, and SM suggest that distributed patterns of structural covariance peak in middle adulthood before returning to a relatively localized topology in older adulthood.

The second significant latent variables of the DN and VAN share spatial features of structural covariance with the second latent variables of the DAN, FPCN, and SM; however, their trajectories (panel F, Figures 2 and 6) do not show a reliable, inverted U-shape. One possible explanation for this is that the selected seed regions for the DN and VAN included regions such as the medial prefrontal cortex, posterior cingulate, and insular cortex. These regions are known functional hubs (van den Heuvel & Sporns, 2013) and, in the second latent variable of DAN, FPCN, and SM, their structural covariance reliably differentiates young adulthood from other portions of the lifespan. In PLS, successive latent variables contribute unique, additional portions of variance. Since these seed regions strongly contribute to the structural covariance of the DN and VAN first latent variables, it is possible that the appearance of a near linear decline in the second latent variable—rather than an inverted U-shaped trajectory—is due to the exclusion of medial prefrontal cortex, posterior cingulate, and insular cortex from the second latent variables of the DN and VAN and their explained covariance. This would suggest that these regions are particularly important in shaping age-dependent patterns of structural covariance.

Previous investigations of structural covariance have found variation in the extent to which networks show age-related changes. For example, relatively flat patterns of structural covariance across adulthood have been seen in the visual network (Li et al., 2013) as well as in temporal, auditory, and cerebellar networks (Hafkemeijer, et al. 2014). Our finding that the visual network did not have a significant second latent variable suggests that there is not a significant age-dependent pattern of structural covariance for this network, in agreement with this previous work.

Contrary to our initial hypotheses, we did find age-dependent structural covariance trajectories for SM, where existing literature suggests that there are little to no age-related changes (Li et al., 2013). In the present work, we find SM to exhibit the same age-dependent pattern of structural covariance as DAN and FPCN. Future longitudinal studies of structural covariance patterns will be important to address the impact of age on specific cortical networks.

Overall, our results therefore suggest that the structural covariance patterns of large-scale neurocognitive networks each have a unique spatial topology; however, neurocognitive networks also show overlapping patterns of age-dependent structural covariance.

### 4.3 Relationship of Structural Covariance to Function

Structural covariance networks have been extensively linked to neural function via their marked disruptions in pathology and pathological aging (Bassett et al., 2008; Hafkemeijer et al., 2016; Spreng & Turner, 2013; Valk, Martino, Milham, & Bernhardt, 2015). Alongside functional connectivity, shared structural covariance has been suggested as a defining characteristic of large-scale networks (Seeley, Crawford, Zhou, Miller, & Grecius, 2009, see also Di et al., 2017). It is worth considering, therefore, these lifespan patterns of structural covariance in light of the existing literature on the development of functional connectivity across the lifespan.

In our work, the first significant latent variable seen in all examined networks showed a stable pattern of structural covariance whose integrity declined across the lifespan. This is similar to patterns of decreasing within network functional connectivity with advancing age (Betzel et al., 2014). The second latent variable seen in all networks—with the exception of the visual network—showed an age-dependent pattern that distinguished young adulthood from both childhood and advanced aging. These results mirror developmental trajectories commonly reported in functional connectivity studies with increased functional integration across networks in childhood, peak functional segregation between networks in young adulthood, and de-differentiation of network functional connectivity in older adulthood (Collin & van den Heuvel, 2013).

The significant overlap of structural covariance trajectories found in the current investigation and those trajectories reported in the functional connectivity literature suggest that a lifespan perspective may help illuminate the noted relationship between structural covariance and neural function. Directly assessing the relationship between structural covariance and functional connectivity, however, is a topic for future research aided by the collection of multi-modal imaging data in lifespan samples (e.g., Glasser et al., 2016; Nooner et al., 2012).

### 4.4. Methodological Considerations

Although this study was able to leverage the increasing amount of anatomical data available in open-access repositories, it included important methodological considerations related to age group definition, scanner acquisition strength, and motion correction. Although we sought to create cohorts representing neurobiologically meaningful age ranges, this resulted in unequal representation in both sample size and age range considered. Our smallest included age group, Age Group 6 (76-94y) included 134 participants, while our largest age group, Age Group 3 (26-35y) included 472 participants. Although differing sample sizes across groups will invariably yield more variable estimates of group-wise covariance, we note that our estimates have statistical power comparable to the smallest group size considered. At 134 subjects, this is still significantly higher powered than current standards for MRI data collection, particularly in lifespan samples. An additional consideration with our selection of age cohorts is the age range considered in each age group. Age Group 4, defined here as ages 36-59, spans a larger time period than any of the other cohorts considered. This was in large part due to the paucity of openly available data for that cohort, particularly when compared to other cohorts such as younger adulthood. The continuing collection of multi-modal data for lifespan initiatives such as NKI-RS (Nooner et al., 2012), the HCP Lifespan Project (Glasser et al., 2016), and UK Biobank (Miller et al., 2016) will increase the availability of high-quality data to investigate such questions.

Differences in scanner acquisition strength across data sources provide an additional important methodological consideration. Several data sources, including those representing the youngest and oldest subjects, were acquired at 1.5T, while young adults were acquired at 3T. Although it is likely that subtle differences between groups may have been introduced by MR field strength, inspection of individual subject scores from the two included lifespan data sources (OASIS and NKI-RS) indicate that these subjects do not show divergent results from those seen in the age-restricted datasets or from one another. This is suggestive of general agreement in structural covariance trends across scanner field strength, as OASIS was collected at 1.5T while NKI-RS was collected at 3T. Further, we controlled for MR field strength across groups by adjusting individually derived subject scores and found similar results for both raw and corrected subject scores. Future work assessing structural covariance across the lifespan should nonetheless aim to examine scans acquired at the same MR field strength and ideally on the same scanner.

A limitation of the current study is the inability to implement motion correction of structural images. Recent work has shown that head motion may introduce artifacts into anatomical images, affecting automated estimates of structure (Alexander-Bloch et al., 2016; Savalia et al., 2016). Although acquisition of a resting-state scan has been proposed to flag high-motion subjects for exclusion from structural analyses (Alexander-Bloch et al., 2016; Savalia et al., 2016), not all of the datasets utilized also provided at least one resting-state scan for each subject. We therefore caution that estimates of age-group differences may be inflated by uncorrected motion. Each of these methodological considerations can be addressed in future work, as comprehensive samples of participants across the lifespan with both structural and functional imaging become increasingly available.

In this study we utilized open-access, cross-sectional data sources to examine structural covariance patterns of six neurocognitive networks across the lifespan. Using multivariate PLS analysis, we found that all networks exhibited stable patterns of network specific structural covariance, and with the exception of the visual network showed a second, age-dependent pattern of structural covariance that mirrored developmental trends seen in the functional connectivity literature. The present results confirm the utility of structural covariance in defining neurocognitive networks and reveal both shared and network specific trajectories of structural covariance across the lifespan.

## Acknowledgements

This work was supported in part by an Alzheimer’s Association grant (NIRG-14-320049) to R.N.S. **NIH Peds** data used in the preparation of this article were obtained from the NIH Pediatric MRI Data Repository created by the NIH MRI Study of Normal Brain Development. This is a multisite, longitudinal study of typically developing children from ages newborn through young adulthood conducted by the Brain Development Cooperative Group and supported by the National Institute of Child Health and Human Development, the National Institute on Drug Abuse, the National Institute of Mental Health, and the National Institute of Neurological Disorders and Stroke (Contract #s N01-HD02-3343, N01-MH9-0002, and N01-NS-9-2314, -2315, -2316, -2317, -2319 and -2320). A listing of the participating sites and a complete listing of the study investigators can be found at http://pediatricmri.nih.gov/nihpd/info/participating_centers.html. **HCP** data were provided by the Human Connectome Project, WU-Minn Consortium (Principal Investigators: David Van Essen and Kamil Ugurbil; 1U54MH091657) funded by the 16 NIH Institutes and Centers that support the NIH Blueprint for Neuroscience Research; and by the McDonnell Center for Systems Neuroscience at Washington University. **NKI** data were obtained from the Nathan Kline Institute – Rockland Sample, Release 5. Principal support for the enhanced NKI-RS project is provided by the NIMH BRAINS R01MH094639-01 (Principal Investigator: Michael Milham). Funding for key personnel also provided in part by the New York State Office of Mental Health and Research Foundation for Mental Hygiene. Funding for the decompression and augmentation of administrative and phenotypic protocols provided by a grant from the Child Mind Institute (1FDN2012-1). Additional personnel support provided by the Center for the Developing Brain at the Child Mind Institute, as well as NIMH R01MH081218, R01MH083246, and R21MH084126. Project support also provided by the NKI Center for Advanced Brain Imaging (CABI), the Brain Research Foundation, and the Stavros Niarchos Foundation. **OASIS** data were supported by the following grants: P50 AG05681, P01 AG03991, R01 AG021910, P50 MH071616, U24 RR021382, R01 MH56584. **ADNI** data used in preparation of this article were obtained from the Alzheimer’s Disease Neuroimaging Initiative (ADNI) database (adni.loni.usc.edu). As such, the investigators within the ADNI contributed to the design and implementation of ADNI and/or provided data but did not participate in analysis or writing of this report. A complete listing of ADNI investigators can be found at: http://adni.loni.usc.edu/wp content/uploads/how_to_apply/ADNI_Acknowledgement_List.pdf. Data collection and sharing for this project was funded by the Alzheimer’s Disease Data collection and sharing for this project was funded by the Alzheimer’s Disease Neuroimaging Initiative (ADNI) (National Institutes of Health Grant U01 AG024904) and DOD ADNI (Department of Defense award number W81XWH-12-2-0012). ADNI is funded by the National Institute on Aging, the National Institute of Biomedical Imaging and Bioengineering, and through generous contributions from the following: AbbVie, Alzheimer’s Association; Alzheimer’s Drug Discovery Foundation; Araclon Biotech; BioClinica, Inc.; Biogen; Bristol-Myers Squibb Company; CereSpir, Inc.; Eisai Inc.; Elan Pharmaceuticals, Inc.; Eli Lilly and Company; EuroImmun; F. Hoffmann-La Roche Ltd and its affiliated company Genentech, Inc.; Fujirebio; GE Healthcare; IXICO Ltd.; Janssen Alzheimer Immunotherapy Research & Development, LLC.; Johnson & Johnson Pharmaceutical Research & Development LLC.; Lumosity; Lundbeck; Merck & Co., Inc.; Meso Scale Diagnostics, LLC.; NeuroRx Research; Neurotrack Technologies; Novartis Pharmaceuticals Corporation; Pfizer Inc.; Piramal Imaging; Servier; Takeda Pharmaceutical Company; and Transition Therapeutics. The Canadian Institutes of Health Research is providing funds to support ADNI clinical sites in Canada. Private sector contributions are facilitated by the Foundation for the National Institutes of Health (www.fnih.org). The grantee organization is the Northern California Institute for Research and Education, and the study is coordinated by the Alzheimer’s Disease Cooperative Study at the University of California, San Diego. ADNI data are disseminated by the Laboratory for Neuro Imaging at the University of Southern California. This manuscript reflects the views of the authors and may not reflect the opinions or views of the NIH.

